# Cluster decomposition-based anomaly detection for rare cell identification in single-cell expression data

**DOI:** 10.1101/2024.02.25.581975

**Authors:** Yunpei Xu, Shaokai Wang, Hong-Dong Li, Qilong Feng, Yaohang Li, Jianxin Wang

## Abstract

Single-cell RNA sequencing (scRNA-seq) technologies have been widely used to characterize cellular landscapes in complex tissues. Large-scale single-cell transcriptomics holds great potential for identifying rare cell types critical to the pathogenesis of diseases and biological processes. Existing methods for identifying rare cell types often rely on one-time clustering using partial or global gene expression. However, these rare cell types may be overlooked in the initial clustering step, making them difficult to distinguish. In this paper, we propose a Cluster decomposition-based Anomaly Detection method (scCAD), which iteratively decomposes clusters based on the most differential signals in each cluster to effectively separate rare cell types and achieve accurate identification. We benchmark scCAD on 25 real-world scRNA-seq datasets, demonstrating its superior performance compared to 10 state-of-the-art methods. In-depth case studies across diverse datasets, including mouse airway, brain, intestine, human pancreas, immunology data, and clear cell renal cell carcinoma, showcase scCAD’s efficiency in identifying rare cell types in complex biological scenarios. Furthermore, scCAD can correct the annotation of rare cell types and identify immune cell subtypes associated with disease, providing new insights into disease progression.

## Introduction

Single-cell RNA sequencing (scRNA-seq) technologies have enabled researchers to analyze gene expression patterns at single-cell resolution^1^, thereby dissecting cellular heterogeneity^2^, while providing new insights into studying the composition and function of cell types within complex tissues^3^. With the advancement of sequencing technology, a larger scale of data becomes available^4^, enabling for not only characterizing the major cell types but also capturing low-frequency cell types^5–7^. These rare cell types exhibit low abundance and have been extensively validated for their significant role in studying the pathogenesis of various diseases, as well as in biological processes such as angiogenesis and immune response mediation, have been extensively validated. For example, circulating tumor cells (CTCs) are indeed very rare in peripheral blood but their metastasis is closely associated with cancer-related death. It is estimated that CTCs account for 1 or fewer cells in every 10^5^ - 10^6^ peripheral blood mononuclear cells (PBMCs)^8^. This limited presence of CTCs poses a substantial challenge to their detection and characterization in cancer research^9,10^. Therefore, in addition to commonly used tools like Seurat^11^ that comprehensively identify major cell types, developing specialized methods to accurately and effectively identify and characterize these rare cell types has become a major challenge in single-cell research.

Prominent algorithms used in recent years for the identification and analysis of rare cell types include finder of rare entities (FiRE)^12^, CellSIUS^13^, ensemble method for simultaneous dimensionality reduction and feature gene extraction (EDGE)^14^, GapClust^15^, GiniClust series methods^16–18^, RaceID series methods^19,20^, SCISSORS^21^, CIARA^22^, and surprisal component analysis (SCA)^23^. These methods identify rare cells from four main perspectives. The first perspective involves proposing a new method for measuring cell rarity in highly variable gene space. FiRE achieves this through an efficient Sketching process that assigns each cell a hash code multiple times, with the number of a hash bucket serving as an indicator of the rareness of its resident cells. It then assigns a consensus rareness score for each cell, identifying cells with scores above a threshold as rare. GapClust identifies rare cell types by examining the variations in Euclidean distance between cells and their 𝑘-nearest neighbors (KNN) in principal component analysis (PCA) transformed subspace. The second perspective focuses on proposing a novel feature selection process. GiniClust introduces a novel gene selection method to identify high Gini genes specific to rare cell types and then uses a density-based clustering algorithm to cluster cells. The CIARA algorithm, based on KNN, identifies potentially rare cells by examining highly locally expressed genes and then applies the Louvain algorithm to cluster with the selected genes. The third perspective is based on clustering results and proposes a novel method tailored to identify rare sub-clusters. CellSIUS identifies candidate marker genes that exhibit a bimodal distribution of expression values within each cluster, then further divides cells into sub-clusters by performing one-dimensional k-means clustering based on the mean expression of each gene set with correlated expression patterns. RaceID identifies outlier cells within each cluster by evaluating the transcript count variability of every gene across all cells and then reassigns each cell to the most highly correlated cluster. SCISSORS employs silhouette scoring for the estimation of heterogeneity of clusters and reveals rare cells in heterogeneous clusters by a multi-step semi-supervised reclustering process. The last perspective involves proposing dimensionality reduction methods for rare cell discrimination, including EDGE and SCA. Furthermore, the integration of multi-omics data has emerged as a promising approach. For example, MarsGT^24^ combines scRNA-seq and scATAC-seq data, using probabilistic heterogeneous graph transformers for rare cell identification.

Although certain successes have been achieved, these algorithms have limitations in terms of both accuracy and robustness. Methods based on highly variable genes may not fully assess whether there are specific signals within these genes that can distinguish rare cell types, making them sensitive to the number of differentially expressed genes. Feature selection-based methods ignore the potential dependence between different genes when identifying rare cell subpopulations. Cluster-based methods may require further analysis of the genes used to distinguish rare types within each cluster. Dimensionality reduction methods may lose important information during processing or be vulnerable to noise and interference, thereby complicating the accurate identification of rare cells. The method integrating multi-omics data needs to account for potential noise from batch effects and other sources of variation^25^, which could complicate the identification of rare cell types.

To overcome these limitations, we propose scCAD, a Cluster decomposition-based Anomaly Detection method to effectively identify rare cell types. Unlike the existing algorithms, scCAD offers an ensemble feature selection method to maximize the preservation of differential signals of rare cell types. During cluster decomposition, scCAD applies iterative clustering based on the most differential signals in each cluster to effectively distinguish rare types or subtypes that are initially challenging to differentiate. Finally, scCAD provides the user with several potentially rare cell clusters.

We benchmark scCAD on twenty-five real scRNA-seq datasets, showcasing its superior capability to identify rare cell types. In the majority of these datasets, scCAD exhibits higher identification accuracy compared to other state-of-the-art methods. In case studies across diverse biological scenarios, including mouse airway, brain, intestine, human peripheral blood mononuclear (PBMC), and pancreas, scCAD accurately identifies rare cell types reported in previous studies, showcasing its robustness and accuracy. In clear cell renal cell carcinoma data, scCAD corrects rare cell annotation mistakes and identifies disease-associated immune cell subtypes, providing novel insights into disease progression. Moreover, the analysis of the results on two large-scale immunology datasets highlights the excellent scalability of scCAD.

## Results

### Overview of scCAD

Single-cell RNA sequencing data often consist of a diverse range of cell types, each with specific functions, and exhibit significant differences in cell counts. This can complicate the identification of rare cell types during initial clustering, as they may be indistinguishable from major cell types based partial or on global gene expression.

To tackle this challenge, scCAD first employs an ensemble feature selection method to effectively retain differentially expressed (DE) genes in rare cell types. Similar to GiniClust and CIARA, scCAD adopts a similar perspective and underscores the importance of the feature selection procedure, which plays a crucial role in clustering. In contrast to traditional approaches that rely solely on the most variable genes for analysis, scCAD combines the most important genes by utilizing initial clustering labels of cells based on global gene expression and a random forest model^26,27^. Then, scCAD proposes an innovative approach by decomposing the major clusters in the initial clustering through iterative clustering based on the most differential signals in each cluster. After cluster decomposition, clusters serve as the fundamental units rather than individual cells. We define the dominant cell type of a cluster as the type to which the majority of cells in the cluster belong. The number of clusters dominated by specific cell types reflects their rarity. The number of clusters dominated by rare cell types is significantly lower than those dominated by major cell types. To enhance computational efficiency, the number of clusters analyzed is reduced by merging some of the nearest clusters. This is accomplished by merging clusters with the closest Euclidean distance between their centers. The set of clusters obtained from the initial clustering, cluster decomposition, and cluster merging are respectively defined as I-clusters (initial clusters), D-clusters (decomposed clusters), and M-clusters (merged clusters). For each cluster in M-clusters, scCAD utilizes differential expression analysis to identify a specific list of candidate DE genes. Due to limited quantity, rare cell types exhibit a higher degree of independence in the corresponding DE gene list of their respective cluster. scCAD employs an isolation forest model^28^ using the candidate DE gene list and calculates the anomaly score of all cells. An independence score is computed by assessing the overlap between highly abnormal cells and cells contained within the cluster, serving as a measure of each cluster’s rarity. Figure 1 shows a schematic pipeline of scCAD, and the Methods section provides a comprehensive explanation of the step-by-step process in scCAD.

**Fig. 1.**
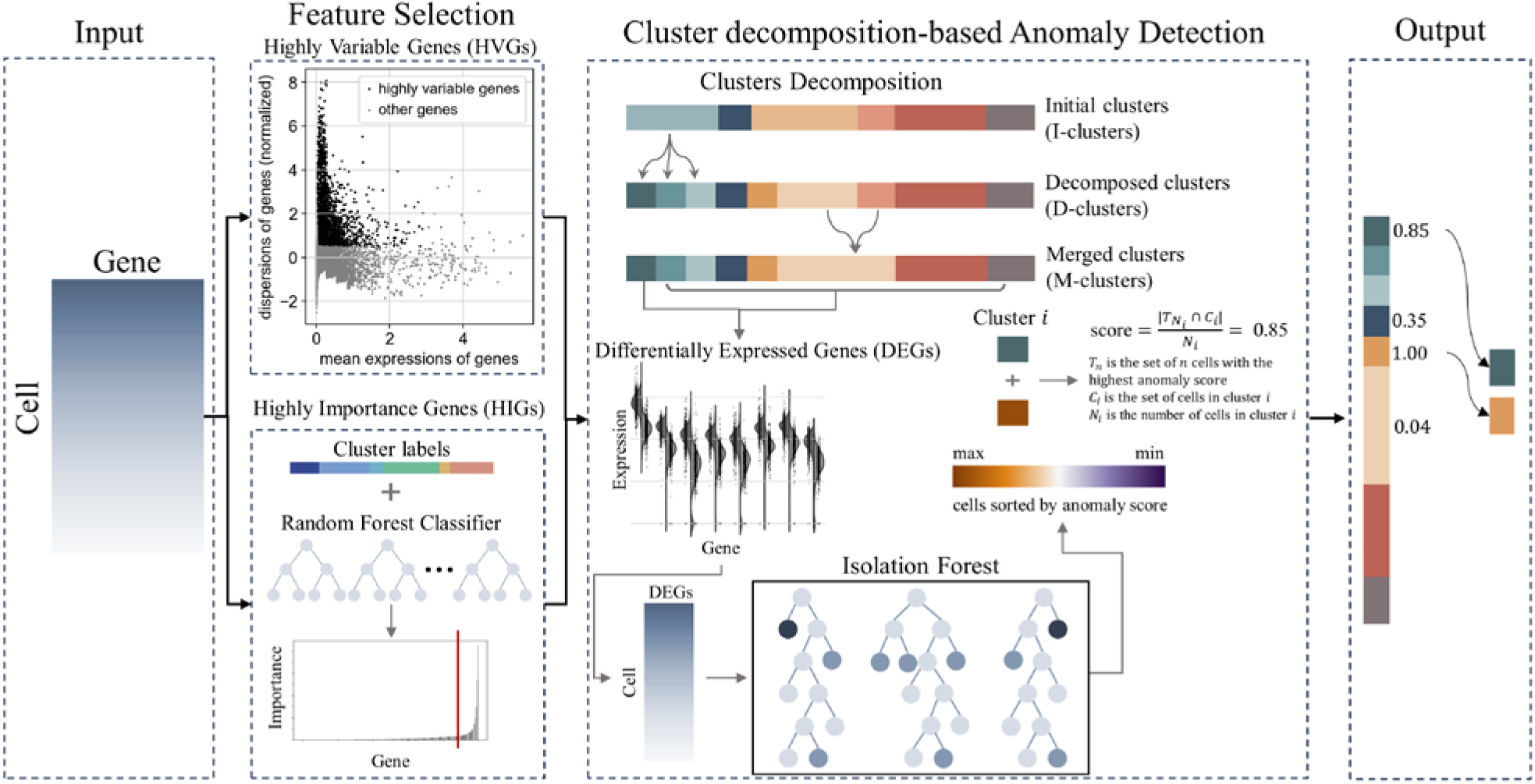
Overview of scCAD. scCAD employs an ensemble feature selection approach, combining the benefits of highly variable genes and highly important genes. It then decomposes the major clusters in I-clusters through iterative clustering. To enhance computational efficiency, certain nearest clusters are merged. For each cluster in M-clusters, scCAD conducts anomaly detection by analyzing the corresponding differentially expressed (DE) genes, assigning an independence score to each cluster. Finally, scCAD provides the user with several potential rare cell clusters according to the independence score.

### Benchmarking scCAD in real datasets

To comprehensively evaluate scCAD, we compare it with ten state-of-the-art methods designed for identifying rare cell types across twenty-five real scRNA-seq datasets representing diverse biological scenarios. The evaluation of different methods is conducted using the F1 score for rare cell types, which effectively captures the trade-off between precision and sensitivity. As shown in Fig. 2a, scCAD achieves the overall highest performance (F1 score = 0.4172) and exhibits performance improvements of 24% and 48% compared to the second and third-ranked methods (SCA: 0.3359, CellSIUS: 0.2812), respectively.

**Fig. 2.**
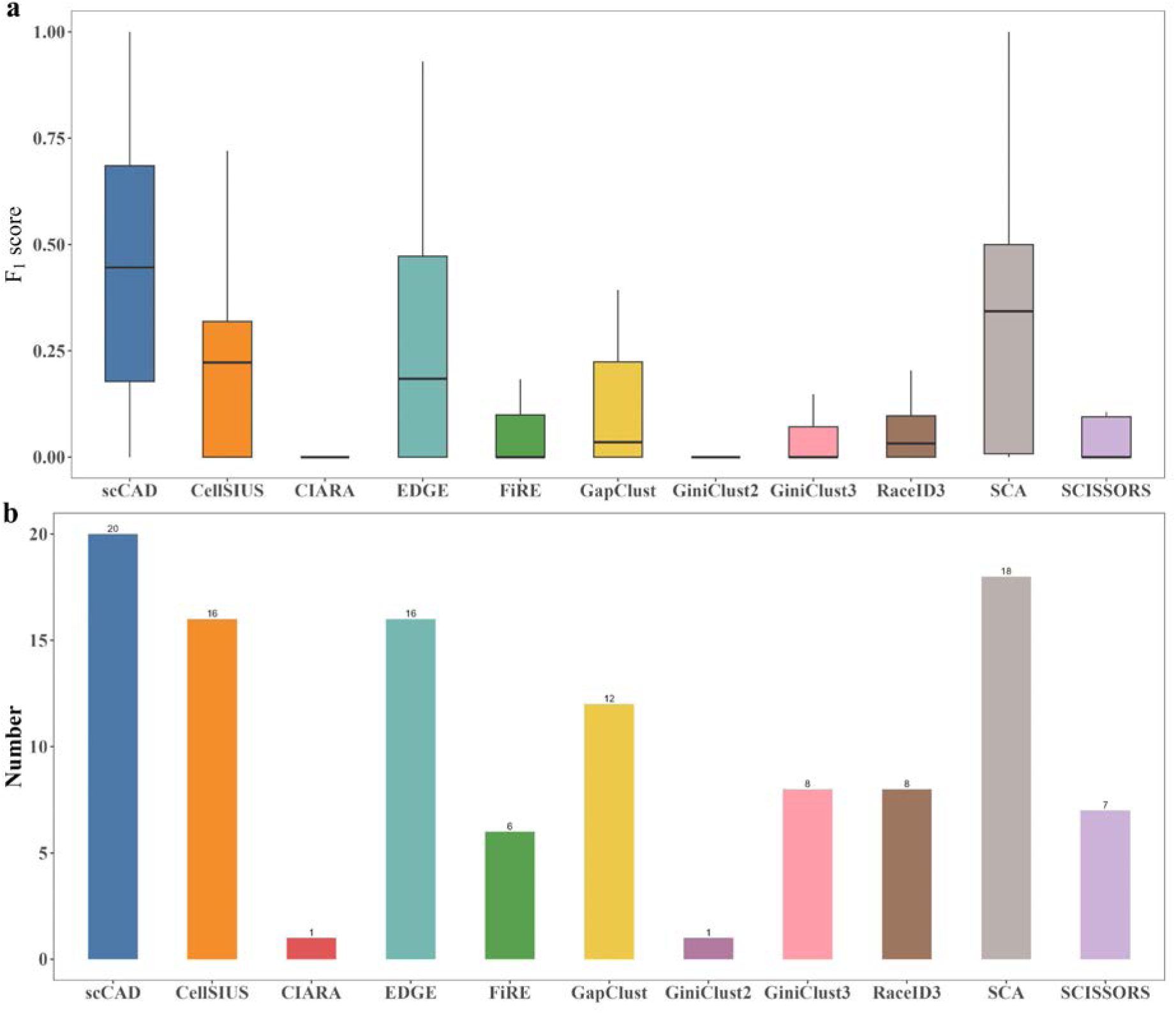
Evaluating scCAD against ten state-of-the-art methods for identifying single-cell rare types on twenty-five real datasets. **(a)** Comparing the distribution of F_1_ scores across all datasets in identifying rare cell types. **(b)** Comparison of the total number of datasets in which at least one rare cell type can be identified successfully.

During the testing process, we observed that several methods are not sufficiently adaptable to all datasets representing diverse scenarios. For example, the series methods of GiniClust may introduce errors by failing to identify high Gini genes^29^. Furthermore, certain methods (such as RaceID) may encounter challenges in generating results for datasets containing more cells due to lower computational efficiency^12^. In contrast, scCAD, EDGE, FiRE, SCA, and SCISSORS can run effectively on all datasets, showcasing their greater suitability for data analysis across a wide range of biological scenarios.

To further evaluate the performance of these algorithms, we count the total number of datasets in which each method successfully identifies rare cell types (Fig. 2b). Since only 30% of the cells are used, it is possible to accurately annotate cell types^30^. Therefore, if a method identifies over 30% of the cells belonging to at least one rare cell type in a dataset, it is deemed to have successfully identified rare cell types in that dataset. As illustrated in Fig. 2b, scCAD demonstrates advantages by successfully identifying rare types in 20 datasets. Meanwhile, we further compare scCAD with four other methods (CellSIUS: 16, EDGE: 16, GapClust: 12, and SCA: 18). We calculate the average F1 score of these five methods on the corresponding datasets where they successfully identified rare cell types. scCAD also demonstrates an advantage (F1 score = 0.5208) compared to the other methods (CellSIUS: 0.3339, EDGE: 0.3954, GapClust: 0.2940, and SCA: 0.4661).

We observe significant variation in the number of rare cell types across different datasets, in which there are 11 datasets with two or more rare cell types and 14 datasets with only one rare cell type. scCAD can identify the rare cell type in, 10 of the 14 datasets and also identify 2 or more rare cell types in 8 of the 11 datasets. In summary, scCAD excels at identifying rare cell types in diverse biological scenarios.

### Feature selection effectively preserves the rare cell type-specific genes

Feature selection is crucial for identifying rare cell types, as it aids in extracting and preserving key features specific to these types, thereby reducing noise and redundant information, and improving the ability to identify and distinguish these types. Most current methods for identifying rare cell types rely on a specific set of highly variable genes (HVG)^12,13,15^, which exhibit significant expression changes across cells, thus potentially providing more information. Our previous study illustrated that highly important genes (HIG) based on random forests have been demonstrated to enhance clustering performance^27^. The gene selection strategy of scCAD involves merging and removing duplicates from the top 2,000 HVG and the top 2,000 HIG. To demonstrate the effectiveness of this strategy, we assess whether the genes selected by scCAD encompass genes specific to rare cell types. Specifically, we first apply Wilcoxon’s rank sum test to identify the top 50 DE genes for each rare cell type in the dataset, which are commonly utilized to indicate the type’s differential signals^31,32^. Then we collect these genes to form a reference gene set 𝑆_ref_, which is regarded as having rare cell type-specific signals. Assume that 𝑆_select_ is the selected gene set, and the overlap rate between the reference gene set and the selected gene set can be calculated as 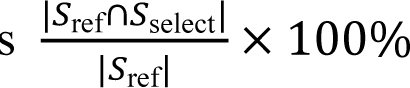. A higher overlap rate indicates a stronger presence of rare cell differences in the selected features. We simultaneously compare scCAD with two individual strategies across all datasets (Fig. 3a). To ensure fairness, the number of features used for HVG and HIG is kept consistent with the number of features finally selected by scCAD. scCAD obtains an average overlap rate of 86.75% in all datasets and exhibits the highest overlap rate in 16 datasets. This indicates that the ensemble feature selection method employed by scCAD can effectively preserve the rare cell type-specific genes.

**Fig. 3.**
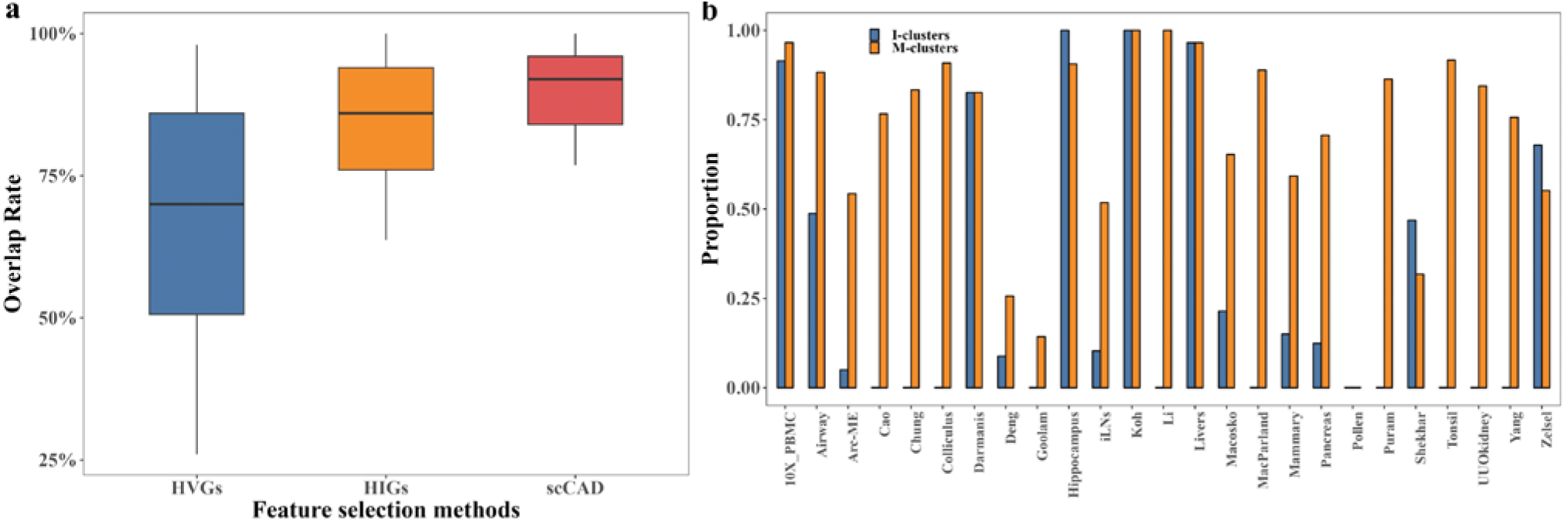
The performance analysis of the feature selection and the cluster merging in scCAD. **(a)** Comparing the distribution of the overlap rate between genes selected by three strategies and the reference differentially expressed genes of rare cell types across all datasets. **(b)** The average proportion of all rare cell types in their dominant clusters is compared between the I-clusters and the M-clusters in all datasets.

### Decomposition effectively isolates clusters dominated by rare cell types

Clusters are commonly annotated based on the primary gene expression patterns of their containing cells, which represent the characteristics of the most dominant cell type. For each cluster, we first count the number of cells of different cell types contained in the cluster based on the annotation information. Then, we identify the cell type with the highest cell count as the dominant type within the cluster. The occupy rate of the dominant cell type in the cluster 𝑖 is defined as follows: 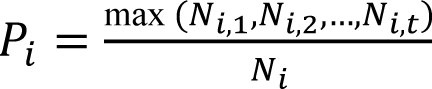, where 𝑁_𝑖,𝑗_ is the number of cells of type 𝑗 in the cluster 𝑖, 𝑡𝑡 is the total number of cell types contained in the cluster 𝑖, and 𝑁_𝑖_ is the total number of cells in the cluster 𝑖. A higher rate serves as an indicator of increased cluster purity, implying that more cells within the cluster belong to the same cell type. For one cell type *j* in one dataset, the proportion of cell type can be calculated as mean(𝑃_𝑗,1_, 𝑃_𝑗,2_, …, 𝑃_𝑗,𝑙𝑗_), where 𝑃_𝑗,𝑥_ is the occupy rate of the cell type 𝑗 in the dominated cluster 𝑥 and 𝑙_𝑗_ is the number of clusters dominated by cell type 𝑗. Subsequently, for each dataset, we separately average proportions of all cell types and rare cell types. To demonstrate the improvement, we compare the average proportions of the clusters from M-clusters with those from I-clusters across all datasets. For a more intuitive representation, we visually present the comparison results of rare types and all types across all datasets (Fig. 3b), respectively.

After cluster decomposition and merging, it becomes evident that the average proportion of cell types within their dominant clusters has significantly increased, especially for rare types, with an average increase from 0.283 to 0.704. Notably, in almost half of the datasets, the initial clustering process fails to identify any rare cell types. In addition, in Fig. 3b, we observe that the average proportions of rare cell types and all cell types in M-clusters are almost higher than those in I-clusters. But we can find from Fig. 3b that neither I-clusters nor M-clusters contain clusters dominated by the unique rare cell type present in the data in the Pollen dataset. The reason for the poor results may be due to the poor separability of this type and almost all methods can not identify the rare cell type in the dataset. Overall, cluster decomposition in scCAD can effectively isolate clusters dominated by rare cell types.

### Evaluation of robustness and sensitivity of scCAD

To analyze the robustness and sensitivity of scCAD with respect to the number of differentially expressed (DE) genes, we conduct tests using an artificial scRNA-seq dataset and a Jurkat scRNA-seq dataset. The artificial scRNA-seq dataset comprises 2,500 cells and two cell types, with the minor cell type representing approximately 1% of the total population. Further details regarding the generation of this dataset can be found in the Methods section. The Jurkat dataset consists of an equal-proportion in vitro mixture of 293T and Jurkat cells^33^. This dataset has been utilized in several previous studies^12–15,34^ to simulate the rare cell phenomenon by adjusting the proportion of Jurkat cells. We generate a subsampled dataset by adjusting the proportion of Jurkat cells to 1%. For both datasets, we set aside the pre-identified differentially expressed (DE) genes which are selected through a stringent criterion, and retain all the non-DE genes in the dataset. Additional details about the identification of DE genes and non-DE genes in both datasets can be found in the Methods section.

Based on the computational efficiency of the algorithm, we compare scCAD with three rare cell detection algorithms: FiRE, GapClust, and GiniClust3. During each iteration of the experiment, an equivalent number of non-DE genes are substituted with randomly selected pre-identified DE genes. This process is repeated 10 times for each number of DE genes. The average F1 score across iterations of different methods is compared for each count of DE genes (Fig. 4).

**Fig. 4.**
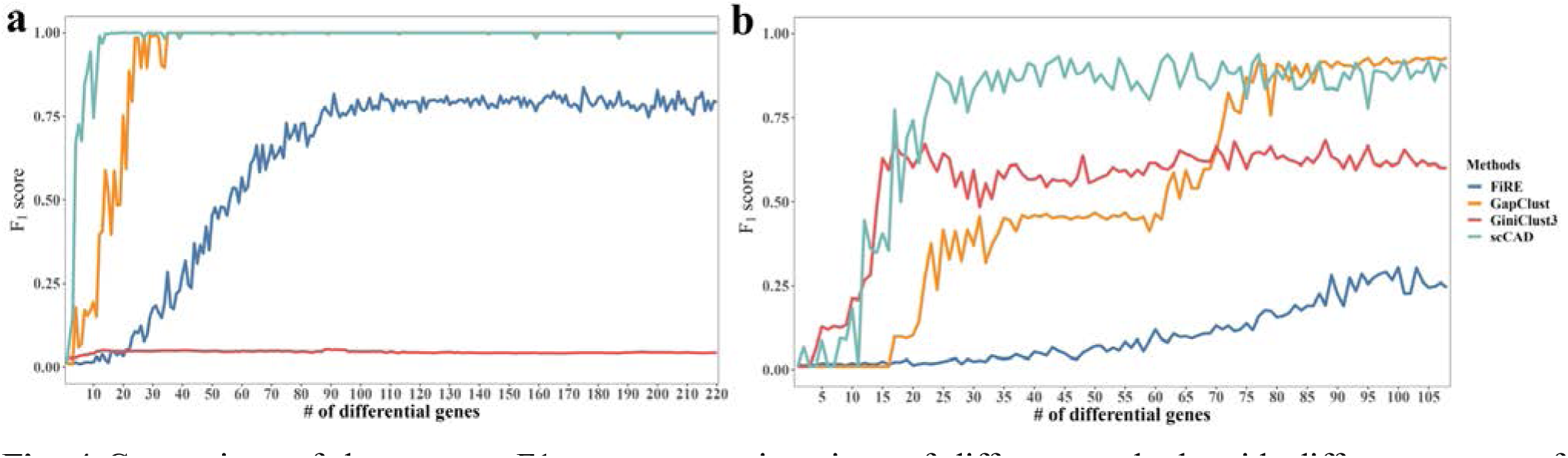
Comparison of the average F1 score across iterations of different methods with different counts of differentially expressed genes on artificial dataset **(a)** and Jurkat dataset **(b)**.

As shown in Fig.4, it becomes evident that scCAD predictions exhibit a notable improvement, maintaining stability at the highest level when 15 and 20 DE genes are introduced, respectively. FiRE and GapClust require more DE genes to achieve a similar stable prediction result. GiniClust3 achieves stability with a similar number of DE genes as scCAD in the Jurkat dataset, but its predictive performance is lower than scCAD. In summary, scCAD demonstrates higher robustness and sensitivity for the number of DE genes.

### scCAD enables the identification of rare airway epithelial cell types

The airways of the lungs are a prominent site for diseases such as asthma, where rare cells play pivotal roles in maintaining airway function^35^. Montoro et al.^36^ utilized scRNA-seq to examine the cellular composition and hierarchy of mouse tracheal epithelium, providing the expression profile of 7,193 cells. They discovered seven cell types, including two rare ones: the Foxi1^+^ lung Ionocyte and Goblet cells. t-distributed Stochastic Neighbor Embedding (t-SNE) is used to visualize the distribution of the cells (Fig. 5a (left)).

**Fig. 5.**
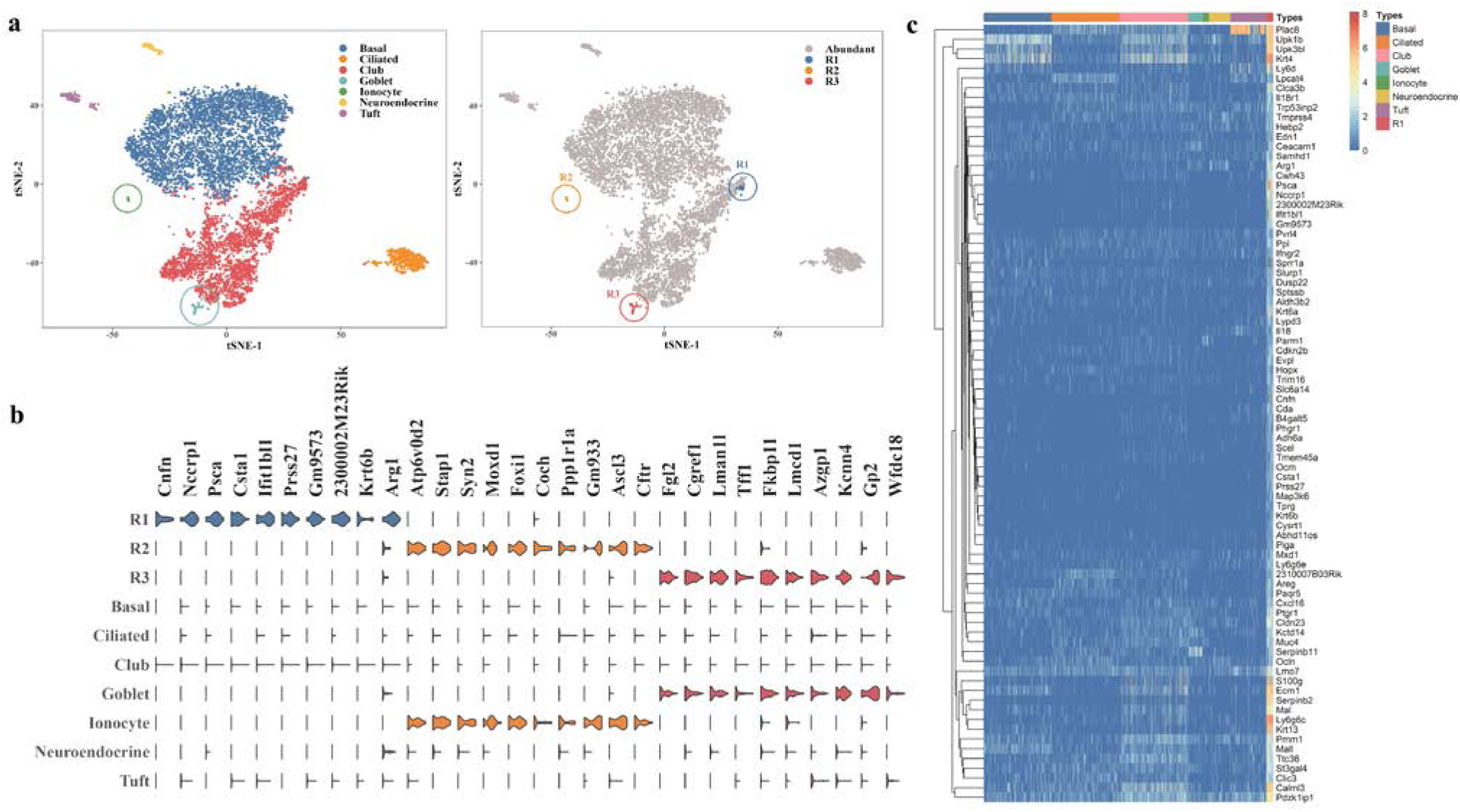
Visualization analysis of scCAD’s results in airway epithelial. **(a)** The t-SNE-based 2D embedding of the cells with color-coded identities (left). Ionocytes and Goblet cells are specifically marked with circles. The three rare cell clusters detected by scCAD are visually distinguished using different colors (right). **(b)** Violin plots showing the expression distribution of the most differentially up-regulated genes in each identified cell cluster. Additionally, seven annotated cell types reported by Montoro et al. are used for comparison. Genes within the same cell cluster are indicated with the same color. **(c)** The expression of all genes differentially up-regulated in cluster R1 is examined across all cell types, including cluster R1 itself.

We apply scCAD to identify rare airway epithelial cell types. scCAD identifies a total of three rare cell clusters, denoted as R1 (0.42%), R2 (0.26%), and R3 (0.57%) (Fig. 5a (right)). We performed differential expression analysis to find cell type-specific marker genes for the newly retrieved cell clusters. Specifically, we use Wilcoxon’s rank sum test to identify differentially up-regulated genes with an FDR cutoff of 0.05 and an inter-group fold-change cutoff of 1.5 for each cluster and each annotated cell type, separately. Assume that 𝑆_𝑖_ is the set consisting of differentially up-regulated genes in identified cluster 𝑖, and 𝑆_𝑗_ is the set consisting of differentially up-regulated genes in annotated cell type 𝑗. The Jaccard similarity coefficient between these two gene sets can be calculated as 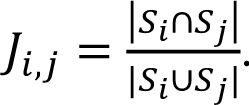. We calculated the similarity between all identified clusters and annotated cell types. For better visualization, we use the top 10 differentially up-regulated genes for each cluster and compare the identified rare cell clusters with annotated cell types based on the expression distribution of these genes (Fig. 5b). As shown in Fig. 5b, clusters R2 and R3 correspond to Ionocytes and Goblet cells, respectively. We discover that cells within the R2 cluster exhibit classic Ionocyte markers, such as the transgenic *Foxi1*-EGFP, the V-ATPase-subunit gene *Atp6v0d2*, the cystic fibrosis transmembrane conductance regulator (*Cftr*) gene, the transcription factor *Ascl3*, and *Smbd1* (formerly known as *Gm933*)^36,37^. Cells within the R3 cluster exhibit classic markers associated with Goblet-1, a subset of Goblet cells as given in Ref. 34. This cluster is enriched for the expression of genes encoding the key mucosal protein (*Tff1*) and secretory regulator (e.g., *Lman1l*).

In contrast to the other two clusters, cluster R1, which consists of 30 cells annotated as Club cells, does not have a corresponding annotated cell type. We visualize the expression of all genes that are specifically up-regulated in cluster R1 across both cluster R1 and all other cell types (Fig. 5c). As shown in Fig. 5c, these genes do not show significant expression in other cell types. Interestingly, we note that R1 shares striking similarities with the “hillock” cells identified by Montoro et al. in their analysis of cell differentiation trajectories. These rare transitional cells connect Basal to Club cells through the unique expression of *Krt13* and *Krt4*^38^. Deprez et al. described a population of *KRT13*^+^ cells in the turbinates, indicating that hillock cells may also exist in other regions of the human respiratory tract^39,40^.

### scCAD identifies various rare cell subpopulations within the mouse brain

In general, the identification of rare cell types becomes more challenging as the dataset encompasses a larger number of cell types, particularly in datasets with multiple cell subtypes^7^. To demonstrate the effectiveness of scCAD in identifying rare cell subtypes in such datasets, we utilize an existing scRNA-seq dataset including 20,921 cells located in and around the hypothalamic arcuate-median eminence complex (Arc-ME)^41^. This dataset encompasses 36 cell subtypes, with 20 of them being considered rare cell subtypes, accounting for proportions ranging from 0.038% to 0.884%. t-SNE is applied to visualize the distribution of the cells (Fig. 6a). For a more intuitive comparison, the cells belonging to rare cell subtypes are color-coded to represent their respective identities in the t-SNE-based 2D embedding (Fig. 6b (left)). scCAD identifies a total of seven rare cell clusters, denoted as R1 (0.87%), R2 (0.63%), R3 (0.40%), R4 (0.50%), R5 (0.12%), R6 (0.11%), and R7 (0.17%) (Fig. 6b (right)). Due to the small number of significantly differentially expressed genes identified in this dataset, we utilize all differentially expressed genes rather than just the up-regulated ones. For better visualization, we select the top 3 differentially expressed genes for each cluster and compare the identified rare cell clusters with annotated cell subtypes based on the expression distribution of these genes (Fig. 6c). As shown in Fig. 6c, cluster R1∼R7 identified by scCAD are highly similar to the seven minor cell subtypes reported by the original study, respectively. Among them, cells within the R1 cluster exhibit gene expression patterns similar to the rare cell subtype annotated as s27.oligodendrocyte6^41^. We discovered that the expression of several characteristic markers in R1 is associated with a subtype of oligodendrocyte known as NFO (newly formed oligodendrocytes)^42^. NFO represents a distinct stage of oligodendrocyte differentiation. Cluster R1 shows characteristic markers including *Fyn*. Additionally, it shows high expression of *Gpr17*^43^, which is involved in oligodendrocyte differentiation, and epigenetic factors such as *Sirt2*, which are also highly transcribed in NFO. Clusters R2 and R7, which include markers such as *Dcn*, *Sparc*, and *Igfbp7*^44–46^, show a high degree of similarity to two distinct fibroblast subtypes. Cluster R3 shows a high degree of similarity to Astrocytes, including markers *Sparcl1*, *Slc1a3*, *Slc1a2*, *Slc6a11*, *Glul*, and *Apoe*^47,48^. Cluster R4 shows a high degree of similarity to a sub-type of pars tuberalis type 1C, including marker *Cyp2f2*. Cluster R5 shows a high degree of similarity to mural cells, including markers myosin light polypeptide 9 regulatory (*Myl9*), and myosin light polypeptide kinase (*Mylk*)^49^. Cluster R6 closely matches a subtype of neurons from the retrochiasmatic area that highly expresses the *Oxt* gene.

**Fig. 6.**
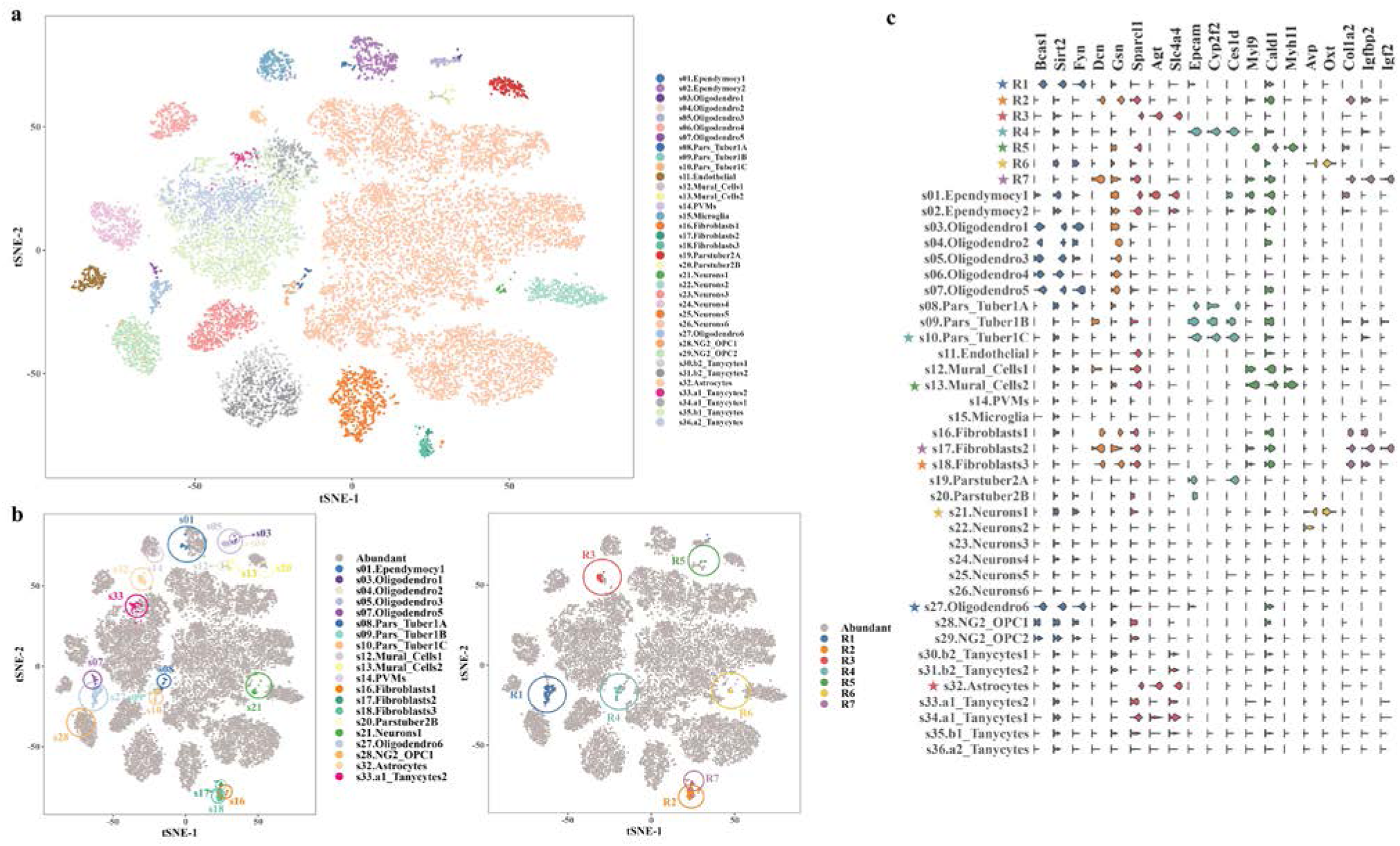
Visualization analysis of scCAD’s results in mouse brain. **(a)** The t-SNE-based 2D embedding of the cells with color-coded identities. **(b)** The t-SNE-based 2D embedding of the cells. The cells in rare cell subtypes are color-coded to indicate their identities (left). The seven rare cell clusters identified by scCAD are visually distinguished using different colors (right). **(c)** Violin plots showing the expression distribution of the most differentially expressed genes in each identified cell cluster. Additionally, 36 annotated cell types reported by Campbell et al are used for comparison. Genes within the same cell cluster are indicated with the same color. Cell clusters that have been identified, along with their corresponding cell subtypes, are marked with an asterisk of the same color.

### scCAD identifies various rare cell types in the crypts of the irradiated mouse intestine

The intestinal epithelium contains various rare cell types, including tuft cells and enteroendocrine cells^50^. Ayyaz et al. conducted scRNA-seq to profile the regenerating mouse intestine and discovered a distinct quiescent cell type called revival stem cell (revSC)^51^, which is induced by tissue damage. Whether it is possible to concurrently detect rare cell types, radiation-induced cell types, and revSCs in enriched crypts after irradiation (IR) is an interesting problem. To solve this problem, we utilize scCAD to analyze an existing scRNA-seq dataset containing 6,644 single-cell transcriptomes of isolated crypts^51^. Ayyaz et al. reported a total of 19 cell clusters. Among them, the 9^th^ and 10^th^ clusters correspond to Enteroendocrine cells, the 18^th^ cluster corresponds to newly discovered revSC, and the 19^th^ cluster corresponds to Tuft cells.

scCAD identifies a total of six rare cell clusters, denoted as R1 (0.90%), R2 (0.50%), R3 (0.63%), R4 (0.56%), R5 (0.21%), and R6 (0.69%) (Fig. 7a). Ayyaz et al. did not annotate the real cell types in this dataset and only provided the most differentially expressed genes for each cluster they reported. Therefore, we calculate the Jaccard similarity coefficient between the set of differentially expressed genes for each identified cell cluster (R1∼R6) and each reported cluster (Cluster1∼Cluster19) as provided by Ayyaz et al. For better visualization, we use the top 10 differentially up-regulated genes for each reported cluster and compare the identified rare cell clusters with reported clusters on the expression distribution of these genes (Fig. 7b).

**Fig. 7.**
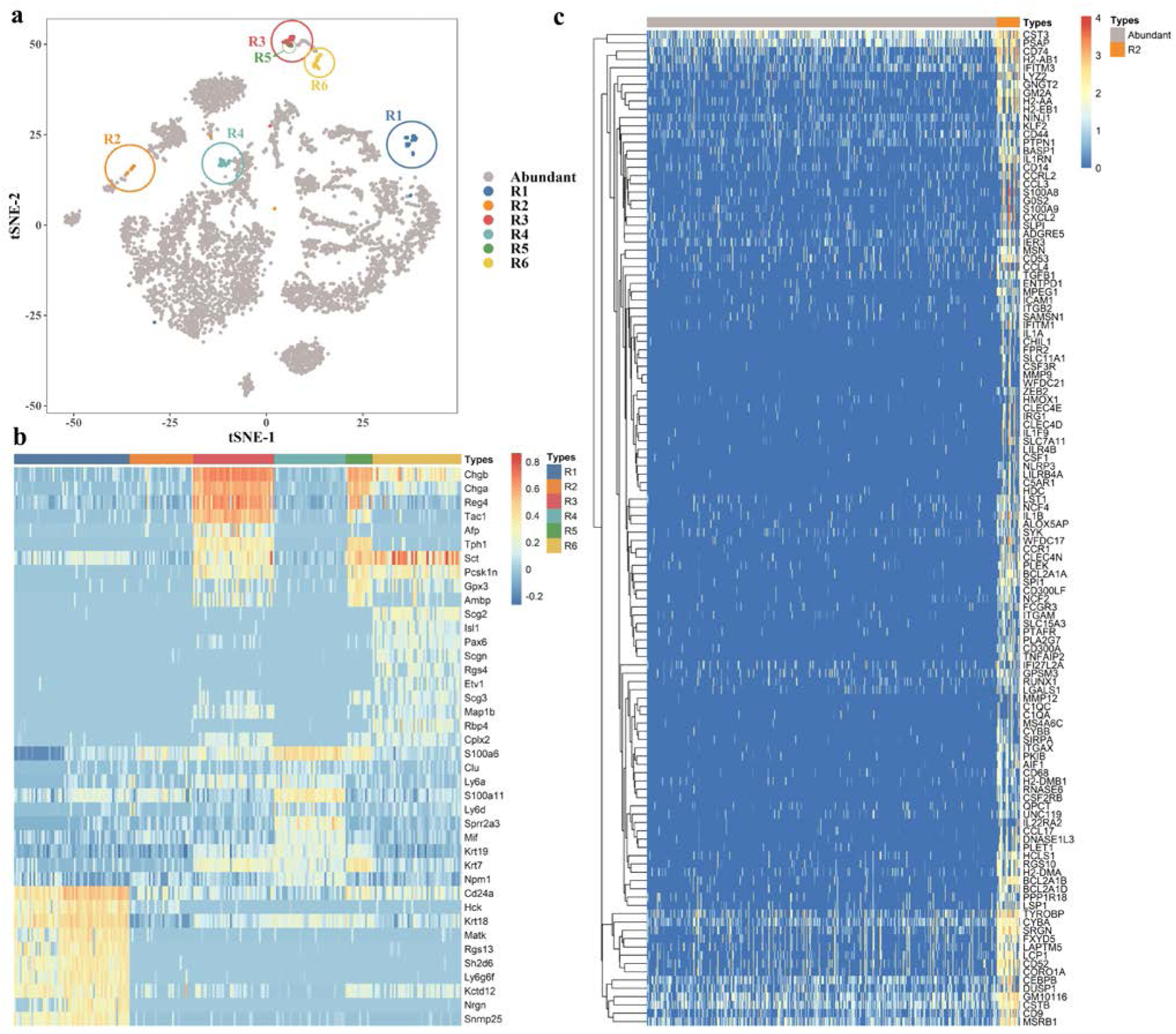
Visualization analysis of scCAD’s results in mouse intestine. **(a)** The t-SNE-based 2D embedding of the cells. The rare cell clusters identified by scCAD are visually distinguished using six distinct colors. **(b)** Expression of the top 10 differentially up-regulated genes from four reported clusters in the cell clusters identified by scCAD. **(c)** The expression of differentially up-regulated genes in cluster R2 across all cells.

As shown in Fig. 7b, we find that Cluster R3, R5, and R6 identified by scCAD are similar to the 9^th^ and 10^th^ clusters, corresponding to Enteroendocrine cells. Clusters R1 and R4 are similar to the 19^th^ and 18^th^ clusters, corresponding to Tuft cells and revSC, respectively. We discover that corresponding cell types markers, such as *Dclk1*, *Trpm5*, *Rgs13*^52^, and *Chga*^53^, are differentially up-regulated in cells within the R1, R3, R5, and R6 clusters. Cluster R4 exhibits gene expression characteristics similar to revSCs, indicating its potential classification as a rare subtype. Notably, we cannot find any clusters reported by Ayyaz et al. that are similar to cluster R2. Consequently, we conduct a more in-depth analysis of the expression of differentially up-regulated genes in cluster R2 across all cells (Fig. 7c).

As shown in Fig. 7c, these genes do not show significant expression in other cells. By querying the PanglaoDB^54^ database for cell type markers, we get that a substantial portion (21%) of the differentially up-regulated genes in cluster R2 corresponded to macrophage markers, including *CD14* and *CD68*^55^. Given the potential association of these rare macrophages with radiation exposure, we conduct additional analysis on other differentially expressed genes and identify *NCF2*, *NCF4*, *CYBB*, and *CYBA* among them. These genes have been observed to exhibit differential expression in the lungs of mice following exposure to IR^56^. They play a crucial role in macrophage activation and polarization towards the M2 subtype. Furthermore, the presence of these macrophages indicates alterations in the inflammatory profile of the irradiated lung tissue^57^.

### scCAD identifies various rare cell types in the human pancreas

The human pancreas comprises various rare cell types such as Epsilon cells^58^. To evaluate the performance of scCAD, we conduct tests on a dataset of 8,569 cells from the human pancreas^59^. This dataset encompasses 14 cell types, with 5 of them being considered rare cell types, accounting for proportions ranging from 0.082% to 0.642%. t-SNE is applied to visualize the distribution of the cells (Fig. 8a (top)). The cells belonging to rare cell types are color-coded to represent their respective identities (Fig. 8a (middle)). scCAD identifies a total of four rare cell clusters, denoted as R1 (0.56%), R2 (0.16%), R3 (0.18%), and R4 (0.33%) (Fig. 8a (bottom)). For better visualization, we select the top 10 significantly differentially up-regulated genes for each cluster and compare the identified rare cell clusters with annotated cell types based on the expression distribution of these genes (Fig. 8b). As shown in Fig. 8b, we find that clusters R2, R3, and R4 correspond to Epsilon cells, Schwann cells, and Mast cells, respectively. We identified distinctive markers associated with these rare cell types within the differentially up-regulated genes, including *GHRL*^60^, *NGFR*, *SOX10*^61^, *KIT*, and *HDC*^62^. In contrast to these clusters, cluster R1, which consists of 48 cells annotated as Beta cells, does not have a corresponding annotated cell type. By further examination, we observed differential up-regulated genes within R1 and compared these cells with other cells belonging to the Beta cell type (Fig. 8c).

**Fig. 8.**
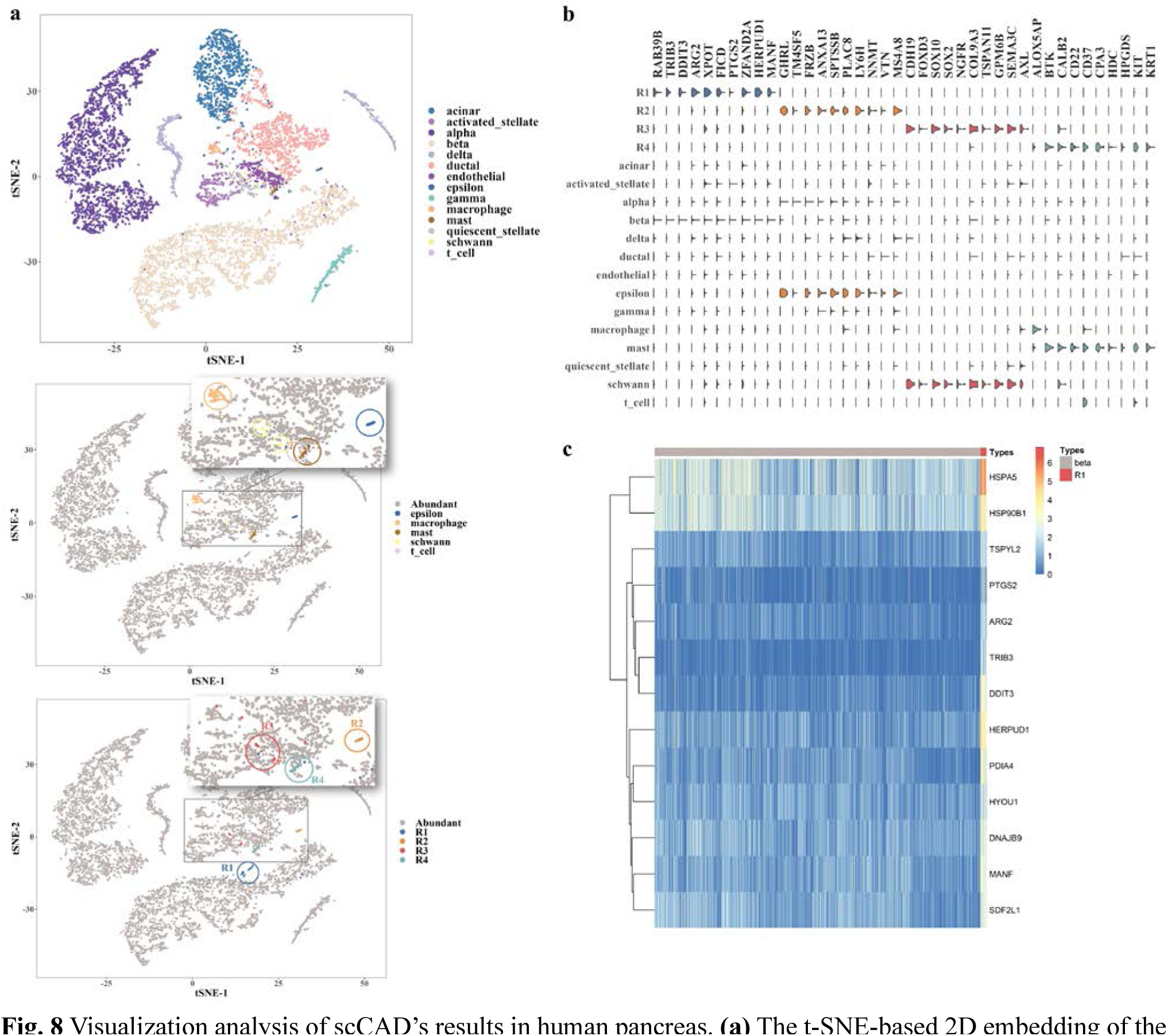
Visualization analysis of scCAD’s results in human pancreas. **(a)** The t-SNE-based 2D embedding of the cells is presented, with color-coded identities indicating cell types (top). Cells belonging to rare cell types are also color-coded (middle). The rare cell clusters identified by scCAD are visually distinguished using four distinct colors (bottom). **(b)** Violin plots showing the expression distribution of the most differentially expressed genes for the four identified cell clusters. Genes within the same cell cluster are indicated with the same color. **(c)** The expression of differentially up-regulated genes in beta cells and cells belonging to cluster R1.

As shown in Fig. 8c, these genes do not show significant expression in other Beta cells. We find that the cells in R1 represent a variant of the Beta cells described by Baron et al.^59^. This variant is characterized by variable expression of genes associated with Beta cell function, such as *HERPUD1*, *HSPA5*, and *DDIT3*^63^, which are involved in endoplasmic reticulum stress response.

### scCAD can identify known rare cell types in large-scale immunological single-cell datasets

To assess scCAD’s ability to detect rare cell types and subtypes in larger single-cell datasets, we collect two immunological datasets separately. One dataset contains 73,259 T cells from 8 human donors^64^, and the other contains 39,563 gastrointestinal immune cells from 10 Crohn’s disease patients^65^. Both of them are well-annotated and comprehensive, making the identification results of scCAD more interpretable. We use t-SNE to visualize cell distribution for both datasets (Fig. 9a), with color-coding cell subtypes to show their identities (Fig. 9b). To visualize the rare cell types in both datasets, we highlight cell types containing less than 1% of the cells in the T cell dataset (Fig. 9b (left)) and immune cell dataset (Fig. 9b (right)).

**Fig. 9.**
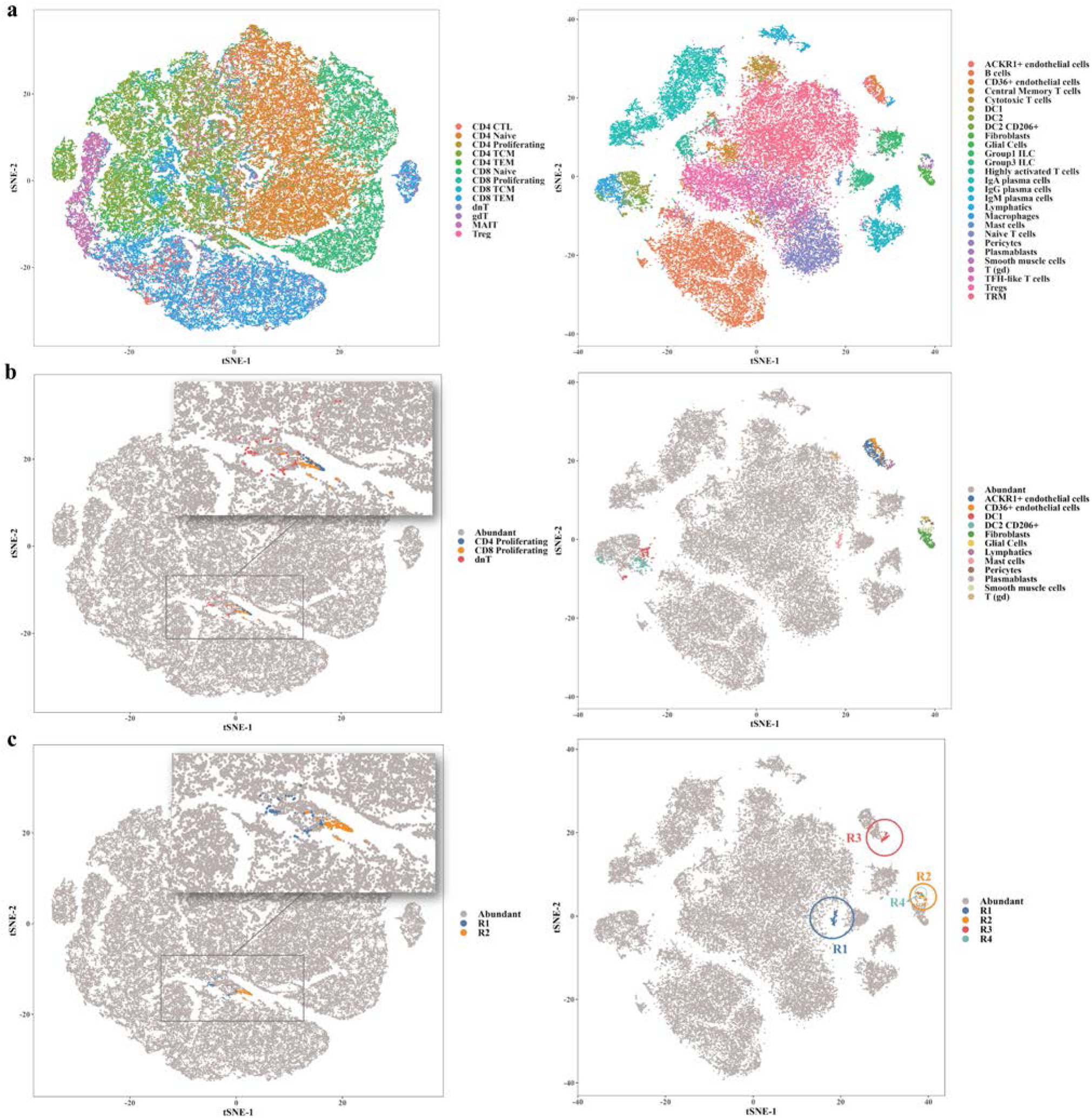
Visualization analysis of scCAD’s results in two large-scale immunological single-cell datasets. **(a)** The t-SNE-based 2D cell embedding with color-coded identities for cell types. The left side depicts the T cell dataset, while the right side shows the immune cell dataset. **(b)** Cell types comprising less than 1% of cells in the T cell dataset are color-coded on the left, while those constituting less than 1% of cells in the immune cell dataset are color-coded on the right. **(c)** The rare cell clusters identified by scCAD are visually distinguished using two and four distinct colors on the T cell dataset (left) and the immune cell dataset (right), respectively.

scCAD identifies two rare cell clusters in the T cell dataset (Fig. 9c (left)): R1 (0.21%) and R2 (0.22%). R1 primarily consists of two types of proliferating cells, CD4 and CD8, with very few annotations in the dataset (0.15% and 0.12% respectively). R1 is mainly composed of double-negative T cells (dnT), which are relatively rare in humans and mice (1∼5% of all T cells)^66^. In the immune cell dataset, scCAD identifies four rare cell clusters (Fig. 9c (right)): R1 (0.29%), R2 (0.26%), R3 (0.27%), and R4 (0.07%). Cluster R1 predominantly consists of mast cells, R2 predominantly consists of pericytes and smooth muscle cells, R3 predominantly consists of lymphocytes, and R4 predominantly consists of glial cells. It’s worth noting that these cell types are the top five rarest annotated in this data.

### scCAD identifies various unannotated rare cell subtypes in the clear cell renal cell carcinoma dataset

Renal cell carcinomas (RCCs) are a diverse group of malignancies believed to originate from kidney tubular epithelial cells. Various RCC subtypes exhibit a broad range of histomorphology, proteogenomic alterations, immune cell infiltration patterns, and clinical behaviors. The most prevalent subtype is clear cell renal cell carcinoma (ccRCC). We collected a total of 6,046 cells annotated into 26 cell clusters from benign adjacent kidney tissues (6 samples from 5 patients) and a total of 20,748 cells annotated into 13 cell types from 7 ccRCC samples^67^. Both of them are utilized to assess the effectiveness of scCAD in the complex tumor microenvironment. As the visualization results of t-SNE are less discriminative for cell types in these two datasets, we visualize the datasets and their respective annotated rare cell types using the 2D Uniform Manifold Approximation and Projection (UMAP)^68^ embedding results (Figure 10a (left), a (middle), b (left), b (middle)).

**Fig. 10.**
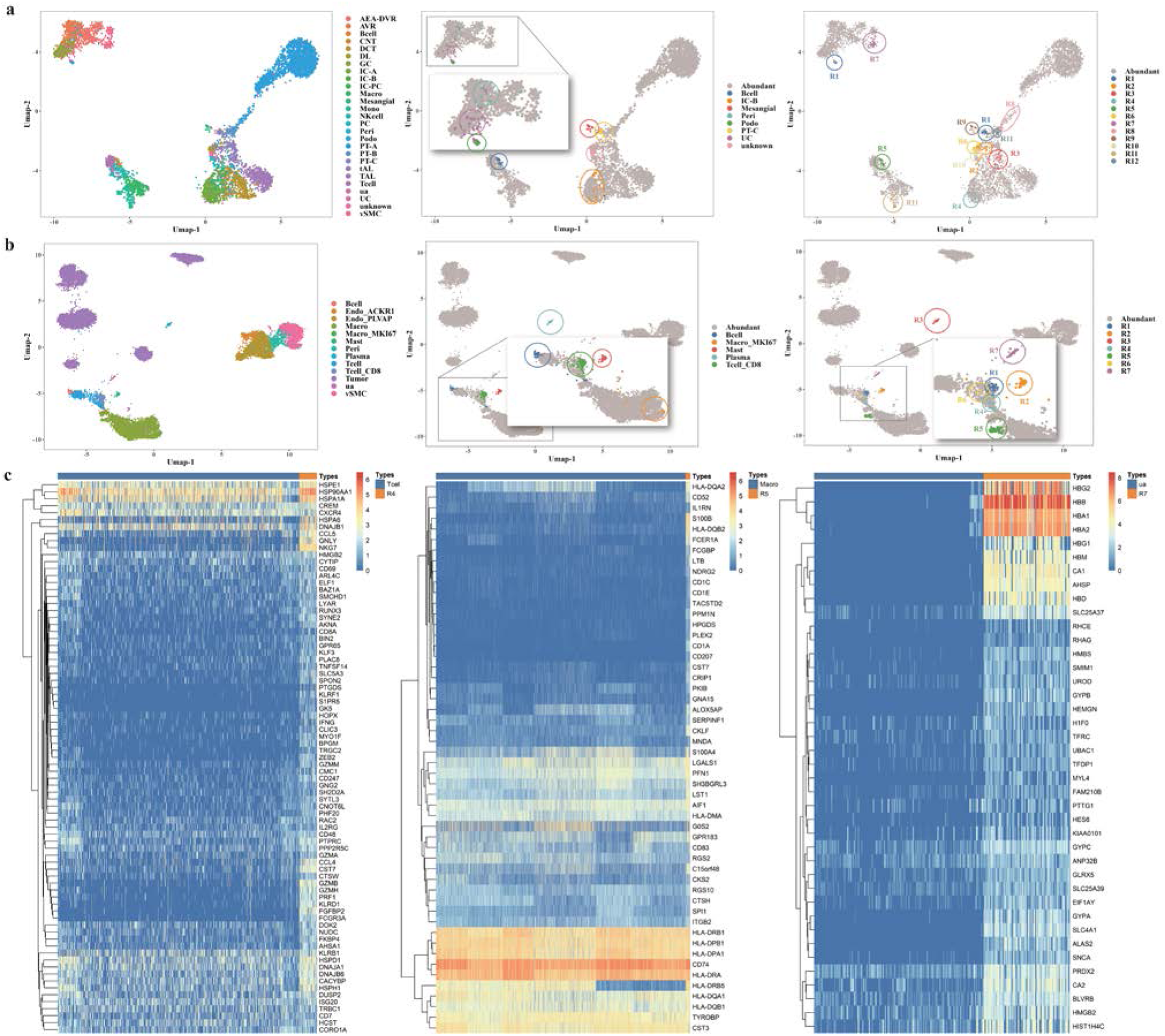
Visualization analysis of scCAD’s results in clear cell renal cell carcinoma dataset. **(a)** UMAP-based 2D visualization depicts cells from the benign kidney, with distinct cell types represented by different color codes (left). Cell types comprising less than 1% of cells are color-coded (middle). The rare cell clusters identified by scCAD are visually distinguished using twelve distinct colors (right). **(b)** UMAP-based 2D visualization depicts cells from the ccRCC, with distinct cell types represented by different color codes (left). Cell types comprising less than 1% of cells are color-coded (middle). The rare cell clusters identified by scCAD are visually distinguished using seven distinct colors (right). **(c)** Comparing the expression of differentially expressed genes in the identified rare cell cluster and other cells annotated as the same type, from left to right: R4 (left), R5 (middle), R7 (right).

In the benign kidney data, scCAD identifies a total of 12 rare cell clusters (0.26%∼0.86%) (Fig. 10a (right)). Upon comparing the detailed annotation information, we discover that the dominant cell types of these clusters encompass multiple rare cell types. For instance, cluster R5 primarily consists of B cells, while R9 is mainly composed of mesangial cells. Notably, scCAD identifies two rare proximal tubule (PT) cell subtypes reported by previous studies^67,69^, namely PT-B (R12) and PT-C (R1).

In the ccRCC data, scCAD identifies a total of 7 cell clusters (0.10%∼0.56%) (Fig. 10b (right)). In addition to CD8^+^ T cells (R1, R6), mast cells (R2), and plasma cells (R3) annotated as rare cell types, scCAD also identifies three rare cell clusters (R4, R5, and R7). Cluster R4 is annotated as T cells. We find that cells in R4 should belong to a rare subtype of effector CD4^+^ T, named CD4^+^ effector-GNLY, characterized by high expression of genes associated with cytotoxicity, including *NKG7*, *GZMB*, *GZMH*, and *GNLY*, as given in a previous study^70^.

Cluster R5 cells are initially annotated as macrophages. However, we identify multiple markers for dendritic cells, such as *CD1C*, *CD207*, and *FCER1A*^71^. Cluster R7 comprises 81 cells from the 239 cells annotated as “ua”. However, we observe that its differentially expressed genes are all related to hemoglobin, including *AHSP*, *HBD*, and *HEMGN*, indicating that this rare cell cluster may be related to hemoglobin synthesis or related biological processes. From the list of highly expressed genes (in reads per kilobase per million transcripts) for each stage of erythroid differentiation^72^, we conclude that cells in cluster R7 are likely in the polychromatic erythroblast stage. Overall, scCAD not only accurately identifies rare cell subtypes but also proves useful in correcting rare cell type annotation mistakes. Furthermore, it has the great potential to identify disease-related immune cell subtypes, providing insights into disease progression.

### Comparative performance of scCAD against multi-omics approach

The advancement in sequencing technology facilitates the integrative analyses of different types of single-cell omics data, providing insights that are more comprehensive than those from a single type of single-cell omics data^73^. This has the potential to enhance downstream analysis performance. However, this progress also presents novel challenges, including the introduction of noise due to batch effects among different omics data^74^. We conduct a comparison between scCAD solely based on scRNA-seq data, and MarsGT^24^, which integrates both scRNA-seq data and single-cell ATAC sequencing (scATAC-seq) data.

Specifically, we first conduct a comparison between scCAD and MarsGT on four real datasets (PBMC-bench-1, 2, 3, and PBMC-test) obtained from human peripheral blood mononuclear cells, which coincide with the datasets used by MarsGT. scCAD solely utilizes the scRNA-seq data in each dataset. We present the performance of scCAD and MarsGT in identifying rare cell types on these datasets, as measured by F1 score, precision, and recall. scCAD demonstrates slightly superior performance compared to MarsGT in terms of F1 score and recall, particularly noticeable in the independent test dataset (PBMC-test), which is the dataset primarily used by MarsGT to illustrate its performance. Upon re-examination of these four datasets, we ascertain that they originate from a common dataset totaling 69,249 cells, with each dataset representing a distinct batch and displaying remarkably similar cell-type distributions. scCAD exhibits greater stability (F1 score standard deviation of 0.1101) compared to MarsGT (0.1747). This difference may result from the effects of technical variations and noise often encountered in the integrative analyses of diverse single-cell omics data types.

Furthermore, we test whether scCAD, using solely scRNA-seq data, could identify rare cell types in the two single-cell multiome datasets employed in MarsGT’s case studies. The two datasets consist of 9,383 cells from the mouse retina^75^ (Retina dataset) and 14,148 cells obtained from a flash-frozen intra-abdominal lymph node tumor (B_lymphoma dataset).

In the Retina dataset, MarsGT reported 12 rare cell clusters, comprising one amacrine cell (AC) cluster, seven bipolar cells (BC) clusters, one horizontal cell (HC) cluster, two Müller glia cell (MG) clusters, and one Rod cell cluster. In contrast, scCAD identifies more rare cell clusters (R1∼R19, totaling 19 clusters). According to the annotations provided in [75], these clusters correspond to two AC clusters (R10, R13), one HC cluster (R8), four Rod cell clusters (R7, R9, R17, R18), six BC clusters (R1, R3, R11, R12, R14, R16), and six retinal ganglion cell (RGC) clusters (R2, R4, R5, R6, R15, R19). Given that BC populations are known to encompass numerous rare populations, we further investigated six clusters associated with BC. We visualize the expression of marker genes specific to the BC subpopulation across the six BC clusters (R1, R3, R11, R12, R14, R16) and the 10 BC subtypes annotated in [75] (Fig. 11a).

**Fig. 11.**
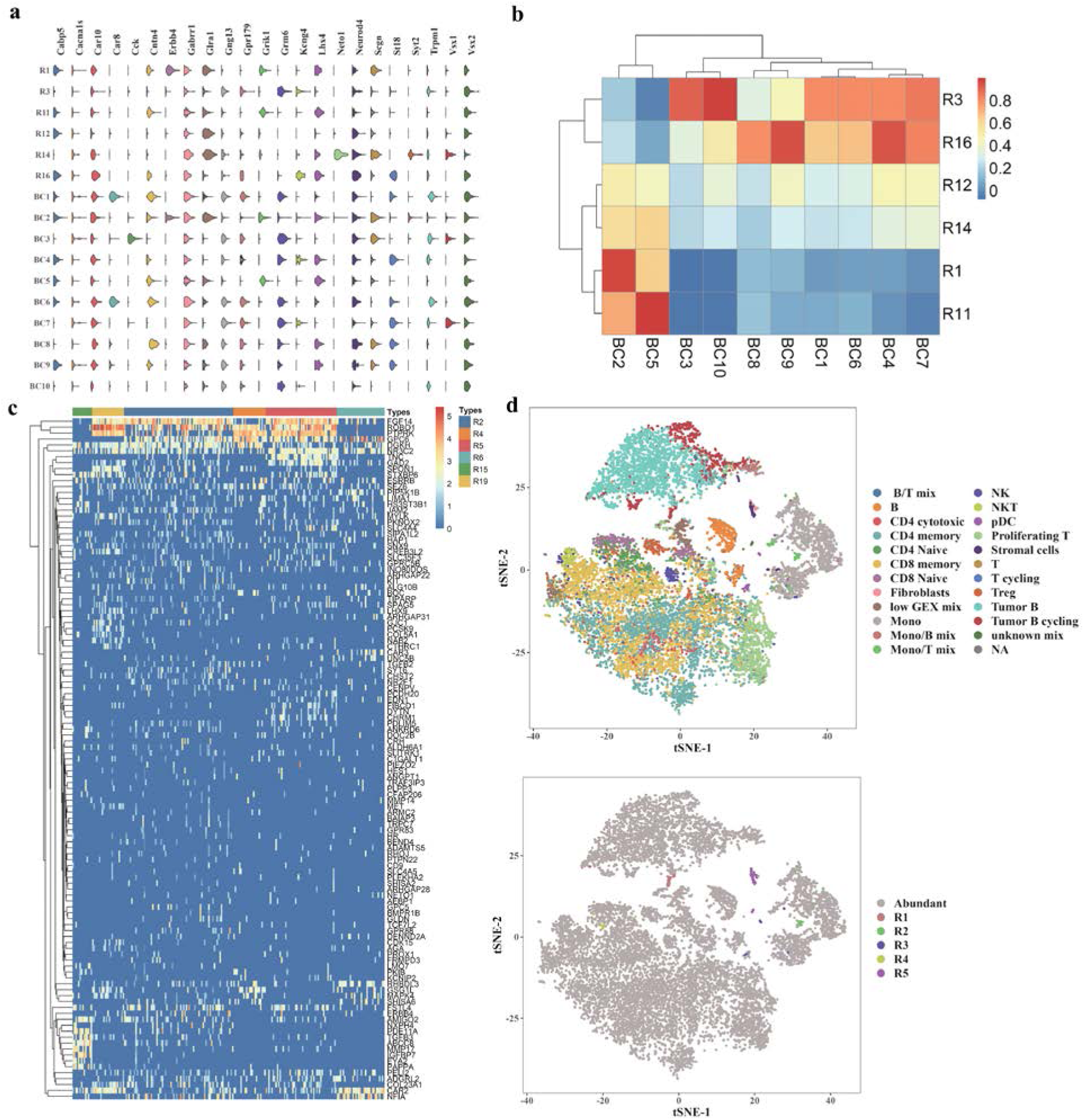
Visualization analysis of scCAD’s results in mouse retina dataset and human lymphoma dataset. **(a)** Violin plots showing the expression distribution of the known marker genes related to BC subtypes across the six identified BC clusters and annotated 10 BC subtypes. **(b)** The Pearson correlation heatmap between the 6 identified BC clusters and the 10 BC subtypes, calculated based on the average expression values of BC marker genes. **(c)** The expression of enriched marker genes from 40 RGC subtypes is examined across all RGC-related clusters (R2, R4, R5, R6, R15, R19). **(d)** The t-SNE-based 2D embedding of the cells with color-coded identities in the lymphoma dataset (top). The five rare cell clusters detected by scCAD are visually distinguished using different colors (bottom).

For better visualization, we compute the Pearson correlation coefficients between the six BC clusters and the 10 BC subtypes based on the average expression values of these marker genes and present them in a heatmap (Fig. 11b). As shown in Fig. 11a and Fig. 11b, we observe that these clusters correspond to distinct BC subtypes, particularly R3, which represents the rarest BC subtype (BC10), accounting for only 3% of all BCs^75^, and was not identified by MarsGT. Additionally, RGCs also exhibit multiple subtypes^76^, prompting us to further analyze the six RGC clusters identified by scCAD. We compile a total of 115 uniquely enriched marker genes from 40 RGC subtypes reported by Rheaume et al.^77^. We visualize the expression of these enriched marker genes across all RGC-related clusters (R2, R4, R5, R6, R15, R19) in Fig. 11c, and we find that these clusters represent various RGC subtypes. Notably, MarsGT only identified a major RGC cluster.

In the B_lymphoma dataset, MarsGT reported a rare state named B lymphoma-state-1. t-SNE is utilized to visualize the cell distribution in the lymphoma dataset (Fig. 11d (top)). scCAD identifies a total of five rare cell clusters (R1∼R5) (Fig. 11d (bottom)). These clusters include one Mono/T mix cluster (R2, 0.27%), one plasmacytoid dendritic cells (pDC) cluster (R3, 0.29%), and one Stromal cell cluster (R5, 0.74%). We identified distinctive markers associated with these rare cell types within the differentially up-regulated genes, including *CD163*^78^, *IL3RA*^79^, and *CALD1*^80^. Unlike the other clusters, neither cluster R1 (0.35%) nor cluster R4 (0.13%) has a corresponding annotated cell type. Through the analysis of their differentially expressed genes, we conclude that they likely correspond to gamma delta T cells and mucosal associated invariant T (MAIT) cells, as indicated by the up-regulated expression of marker genes *CENPF*^81^ and *KLRB1*^82^, respectively. In contrast to scCAD, MarsGT did not identify these rare cell types.

Moreover, scRNA-seq data is more readily accessible, thereby streamlining the data acquisition and processing workflow and reducing experimental costs. In summary, scCAD holds advantages in performance, stability, and cost-effectiveness.

## Discussion

scCAD offers an ensemble feature selection method to maximize the preservation of differential signals of rare cell types, thereby enabling the accurate identification of rare cells. During cluster decomposition, scCAD applies iterative clustering based on the most differential signals within clusters to effectively distinguish rare types or subtypes that are initially challenging to differentiate. With the application of the anomaly detection algorithm, scCAD can identify clusters dominated by rare cell types within the cluster decomposition results. Extensive experiment results show that scCAD demonstrates performance advantages across diverse biological scenarios.

Several computational methods have been developed specifically for identifying rare cell types, broadly categorized into four groups based on their methodological characteristics: feature selection, clustering, dimensionality reduction, and rarity measurement. However, single-cell data often arise from diverse and complex biological scenarios, and the performance of many methods is limited due to imperfect assumptions. In contrast, scCAD adeptly addresses these challenges and offers corresponding solutions.

Indeed, several studies have underscored the pivotal role of feature selection in downstream single-cell data analysis^83–85^. It is commonly acknowledged that the efficient selection of marker genes capable of distinguishing rare cells is crucial for their accurate identification. Extensive testing across various scRNA-seq datasets reveals that highly variable genes often do not fulfill this objective well. However, scCAD successfully identifies over 85% of rare cell-specific expressed genes on average.

Given that single-cell data often contain multiple cell types that differ significantly in number and function, it is difficult for initial clustering to distinguish rare types. To overcome this challenge, scCAD conducts cluster decomposition by iteratively clustering based on the most differential signals in each cluster for the first time. The substantial average proportion of dominant rare types within the clusters after decomposition underscores the effectiveness of this approach in isolating rare cell types. Clusters in M-clusters dominated by rare cell types exhibit stronger rare signals than individual cells. scCAD utilizes an anomaly detection algorithm in the space of cluster-specifically expressed genes. It calculates an independence score based on the overlap of cells in the cluster with highly abnormal cells to determine the rarity. Essentially, the features of rare clusters exhibit higher independence, leading to a clear distinction of cells within them from cells belonging to the major cluster in this feature space. In the benchmark on 25 real-world datasets, scCAD outperforms 10 other state-of-the-art methods in accuracy, successfully identifying rare cell types in 20 of them. In-depth case studies across various complex biological scenarios, including mouse airway epithelium, hypothalamic arcuate median eminence complex (Arc-ME), irradiated mouse intestinal crypts, human pancreas, and large-scale human immunology cells, demonstrate scCAD’s capability in accurately identifying rare cell types and subtypes. This holds even in cases involving radiation and multiple subtypes. Furthermore, in the clear cell renal cell carcinoma (ccRCC) dataset, scCAD not only rectifies the annotation of rare cell types but also identifies rare subtypes not discovered in the original article. Importantly, in diverse datasets containing immune cells, scCAD identifies multiple immune cell subtypes associated with disease, potentially providing new insights into disease progression.

In summary, scCAD has demonstrated its effectiveness as a tool for identifying rare cells, showcasing its high accuracy, sensitivity, and robust generalization capabilities across various biological scenarios.

## Methods

### Data preprocessing

We preprocess the gene expression matrix as follows. First, we filter out genes with low expression rates, which may not provide effective information. Specifically, genes that are expressed in at least three cells are retained for downstream analysis. Each scRNA-seq dataset is normalized by using the log-normalization procedure including the calculation of cell-specific size factor based on the sequencing depths, and normalization. The normalized matrix is then log2-transformed after adding 1 as a pseudo-count.

### Rapid clustering module for single-cell analysis

scCAD utilizes this rapid single-cell clustering module multiple times to assign cluster labels to all or a subset of cells. Similar to previous works^86,87^, scCAD first applies PCA to obtain the top principal components (PCs) that represent the most differential signals in the data. Then, the cell undirected graph is constructed using Euclidean distance and the KNN algorithm, with edge weights uniformly set. Finally, the graph-based community detection algorithm, such as Louvain^88^, is used to assign cluster labels to cells.

### The procedure of scCAD method

scCAD integrates an ensemble feature selection method and a cluster decomposition-based anomaly detection score step. Specifically, scCAD involves the following detailed procedures after data preprocessing.

1. The ensemble feature selection process involves selecting highly variable genes^11^ and highly discriminative genes^27^. Specifically, scCAD calculates the mean and a dispersion measure (variance/mean) for each gene across all single cells, selecting the top 2,000 most variable genes that exhibit high variability compared to genes with similar average expression. At the same time, a random forest model is trained using the preprocessed gene expression matrix and cluster labels, and the importance of each gene is calculated based on the Gini impurity obtained from a set of decision trees^26^. Next, scCAD selects the top 2,000 genes with the highest importance. Finally, the combined set of genes from both selections is retained for subsequent analysis.
2. Using the preprocessed expression matrix with selected genes, scCAD performs cell clustering and initially partitions 𝑛 cells into several clusters. The set of clusters obtained from the initial clustering is defined as I-clusters (initial clusters). Then, scCAD iteratively decomposes each cluster containing more than 𝑅 = 1% of the total number of cells (𝑅 ∗ 𝑛) through clustering until no new clusters are generated by using the Louvain method, or until all clusters become smaller than 𝑅 ∗ 𝑛. The set of clusters obtained from cluster decomposition is defined as D-clusters (decomposed clusters). scCAD merges the nearest neighbor clusters to enhance efficiency and decrease the number of analyzed clusters. Specifically, scCAD first determines the centers for each cluster. The center of cluster 𝑖 is calculated as: *V_i_* = 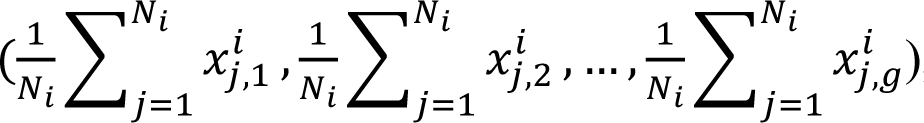, where 𝑉_𝑖_ is a vector with a magnitude of 𝑔, 𝑔 _𝑖_is the number of selected genes, 𝑁_𝑖_ represents the number of cells in cluster 𝑖, and 𝑥^𝑖^ represents the expression value of gene 𝑘 in cell 𝑗 belonging to cluster 𝑖. Then, scCAD calculates the Euclidean distance between all cluster centers: 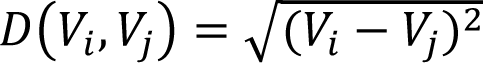, where 𝑉_𝑖_and 𝑉_𝑗_ represent the centers of cluster 𝑖 and 𝑗, respectively. Finally, scCAD determines the threshold of merging, 𝑇 = median(𝑑_1_, 𝑑_2_, …, 𝑑_𝑚_), where 𝑑_𝑖_ is the Euclidean distance between cluster 𝑖 and its nearest neighboring cluster. If 𝐷(𝑉_𝑖_, 𝑉_𝑗_) < 𝑇, clusters 𝑖 and 𝑗 are merged. The set of clusters obtained after cluster merging is defined as M-clusters (Merged clusters): {𝐶_1_, 𝐶_2_, …, 𝐶_𝑚_}.
3. For each gene in cluster 𝑖 in M-clusters, scCAD calculates the difference between the median of the gene expressions of all cells within cluster 𝑖 and the median of those outside of cluster 𝑖^89^. Assume that 𝑋𝑋^𝑘^ represents the vector composed of gene 𝑘 expression values for all cells in _𝑖_ cluster 𝑖, 𝑋𝑋^𝑘^ represents the vector composed of gene 𝑘 expression values for all cells outside _𝑖_ of cluster 𝑖, the median difference of gene 𝑘 is calculated as: 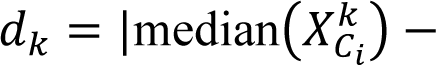 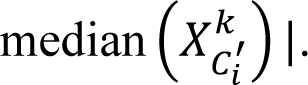. Finally, scCAD selects the top 20 genes with the largest differences to generate _𝑖_ the candidate gene set 𝑆_𝑖_ for cluster 𝑖.
4. The gene expression matrix, which contains the genes in 𝑆_𝑖_, is then fed into an isolation forest model^28^ to calculate an anomaly score for each cell. The isolation forest model builds a collection of isolation trees where each tree is constructed by randomly selecting a subset of cells and recursively partitioning them into smaller subsets based on their expression in randomly selected candidate genes. This process continues until each cell is isolated in its leaf node. The anomaly score is computed by normalizing the average path length for each cell, achieved by comparing it to the average path length of a randomly generated cell from the same dataset. The resulting score represents the degree of abnormality exhibited by each cell with the candidate genes. As described in [28], the ensemble anomaly score of cell 𝑗 based on the candidate genes in 𝑆_𝑖_ is calculated with:

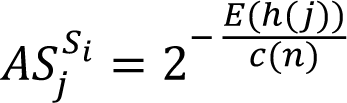

where ℎ(𝑗) is the path length of cell 𝑗 in an isolation tree, which is the number of edges traversed in an isolation tree from the root node to the node containing cell 𝑖. 𝐸(ℎ(𝑗)) is the average of ℎ(𝑗) across all the isolation trees in isolation forest model. 𝑐(𝑛) represents the average path length when the total number of cells is n, and its formula is as follows^28^:

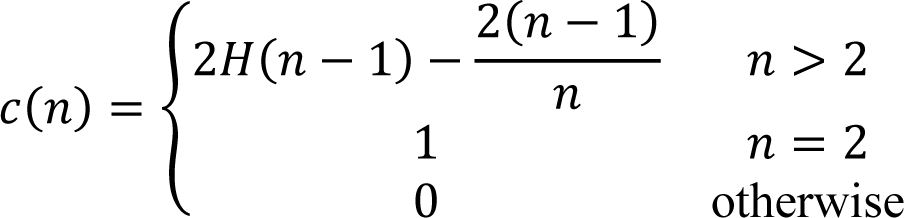

where 𝑇(𝑛 − 1) is the harmonic number that can be estimated by ln(𝑛 − 1) + 0.5772156649 (Euler’s constant)^28^.
5. scCAD assigns an independence score to cluster 𝑖 in M-clusters based on the list composed of the corresponding anomaly scores of all cells: 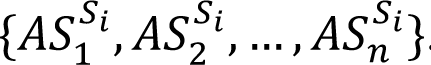. The independence score (IS) 1 2 𝑛 of cluster *i* is defined as follows:

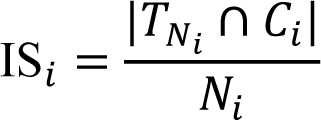

where 𝑁_𝑖_ is the number of cells in cluster 𝑖, 𝑇_𝑁𝑖_ is the set of the top 𝑁_𝑖_ cells with the highest anomaly scores, and 𝐶_𝑖_ is the set of cells in cluster 𝑖. A higher independence score indicates that the differentially expressed genes of the corresponding cluster effectively distinguish and characterize its encompassing cells.
6. scCAD executes steps 3∼5 for each cluster in M-clusters until obtaining the independence score for all clusters: {IS_1_, IS_2_, …, IS_𝑚_}. Finally, clusters with an independence score exceeding the threshold 𝐼 (default is 0.7) ({𝐶_𝑖_ ∈ M-clusters|score_𝑖_ > 𝐼}) are labeled as rare cell types and are outputted, along with the corresponding candidate genes.

### Parameter value selection in scCAD

Clusters containing more than 𝑅 ∗ 𝑛 cells are considered for decomposition through iterative clustering. Based on a comprehensive review of previous studies^18,90,91^ defining the size of rare cell types, we set this parameter to 1% on all larger datasets. For smaller datasets, especially when the total number of cells is below 3,000, we set the threshold to 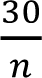 to prevent the generation of excessively small clusters, thereby enhancing the interpretability and reliability of clustering outcomes.

After cluster decomposition, scCAD merges clusters if their distance is smaller than the threshold 𝑇, which is denoted as 𝑇 = median(𝑑_1_, 𝑑_2_, …, 𝑑_𝑚_), where 𝑑_𝑖_ is the Euclidean distance between cluster 𝑖 and its nearest neighboring cluster. We test the number of clusters generated after merging and the average proportions of all cell types and rare cell types in their dominant clusters by using different 𝑇 values in the Arc-ME dataset. Lower 𝑇 values (such as zero and the lower quartile) may incur higher computational overhead due to a larger number of analyzed clusters, while higher 𝑇 values (such as the upper quartile and the 90th percentile) may significantly increase the likelihood of merging clusters dominated by rare cell types, potentially diminishing the effectiveness of the decomposition step. To enhance efficiency and reduce the number of analyzed clusters, we use the median as the default parameter across all datasets. This facilitates the merging of clusters dominated by the same cell types.

scCAD identifies a cluster as rare when its independence score exceeds a threshold value, 𝐼. We display the distribution of independence scores calculated by scCAD for each cluster on four datasets. Clusters dominated by rare cell types exhibit significantly higher independence scores compared to other clusters, and using the default threshold can effectively distinguish them. It is important to note that reducing this threshold may result in the identification of multiple clusters dominated by larger cell types. We default to applying 𝐼 = 0.7 across all datasets.

### Usage of comparative methods

To evaluate the performance of scCAD for identifying rare cells, we conduct a benchmark analysis comparing scCAD with other methods. The CellSIUS package is obtained from GitHub (Novartis/CellSIUS). The initial major cell types are determined using a single-cell clustering workflow in the Seurat package^11^. The CellSIUS algorithm provides the results of sub-clusters assigned to each cell. The CIARA package is obtained from GitHub (ScialdoneLab/CIARA). The CIARA algorithm merges the clustering results obtained by the standard algorithm (Louvain) based on the HVG and identified genes, especially labeling rare cell types. The EDGE package is obtained from GitHub (shawnstat/EDGE). For each dataset, we utilize the data matrix preprocessed by the Seurat package^11^ as the input for this method. Based on the 2-dimensional embedding results generated by EDGE, we construct a 𝑘-nearest neighbor graph and apply the Louvain algorithm to obtain the global clustering results. The FiRE package is obtained from GitHub (princethewinner/FiRE, R version). FiRE assigns a score to each cell and outputs predicted rare cells that meet the thresholding criteria based on the interquartile range (IQR). The GapClust package is obtained from GitHub (fabotao/GapClust). Similar to scCAD, GapClust generates multiple sets of predicted rare cells as output. The GiniClust2 and GiniClust3 packages are obtained from GitHub (dtsoucas/GiniClust2, rdong08/GiniClust3), respectively. Both of them return global consensus clustering results. The RaceID3 package is obtained from GitHub (dgrun/RaceID3_StemID2_package). RaceID3 returns a list containing predicted rare cells. SCA is implemented in Python, and the latest version is obtained from GitHub (bendemeo/shannonca). Following their previous recommendations, we construct a 15-nearest Euclidean neighbor graph in the 50-dimensional space of SCA and use Leiden clustering with the default resolution of 1.0 to obtain the final clustering results. The SCISSORS package is obtained from GitHub (jr-leary7/SCISSORS). We re-clustering the clusters with an average silhouette coefficient calculated by SCISSORS smaller than the overall average, resulting in the final clustering results.

For each algorithm, all parameters are set to their default values. For algorithms that directly output sets of predicted rare cells, such as scCAD, GapClust, FiRE, and RaceID3, we combine all the predicted cells from the result to obtain the final binary prediction outcome. For algorithms that return global clustering results, such as CellSIUS, CIARA, EDGE, GiniClust2, GiniClust3, SCA, and SCISSORS, clusters with a cell population smaller than the corresponding threshold (1% or 5%) are identified as rare clusters. We combine all the cells from the predicted rare clusters to obtain the final binary prediction outcome.

### Generation and description of simulation data

To analyze the sensitivity of scCAD to rare cell-type identity, we generate an artificial scRNA-seq data using the splatter R package^92^. The following command is used to generate these data:

*splatSimulate(group.prob = c(0.99, 0.01), method = ’groups’, verbose = F, batchCells = 2500, de.prob = c(0.4, 0.4), out.prob = 0, de.facLoc = 0.4, de.facScale = 0.8, nGenes = 5000, seed=2023)*

The dataset consists of 2,500 cells, each containing 5,000 genes. Of these cells, 2,476 represent the major cell type, while 24 define the minor type.

After data preprocessing, we use Wilcoxon’s rank sum test to identify DE genes with an FDR cutoff of 0.05 and an inter-group absolute fold-change cutoff of 1.5. Assume that the set of these DE genes is 𝑆_𝐷𝐸𝐷_. We remove the DE genes obtained from the randomly permuted labels from the set 𝑆_𝐷𝐸𝐷_. This step is repeated ten times. The final 220 DE genes are removed from the data and preserved as a separate set. Additionally, 3,226 genes with a p-value exceeding 0.05 are retained as a distinct set of non-differential genes.

The subsampled Jurkat dataset consists of 1,556 cells, each containing 32,738 genes. Of these cells, 1,540 represent the major 293T cell type, while 16 define the minor Jurkat cell type. To increase the number of DE genes for analysis, the inter-group absolute fold-change cutoff is adjusted to 1. The final 108 DE genes are removed from the data and preserved as a separate set. Additionally, 30,479 genes with a p-value exceeding 0.05 are retained as a distinct set of non-differential genes.

## Data availability

All described datasets are obtained from various public websites, including NCBI Gene Expression Omnibus (GEO, https://www.ncbi.nlm.nih.gov/geo/), ArrayExpress (https://www.ebi.ac.uk/arrayexpress/), Sequence Read Archive (SRA, https://www.ncbi.nlm.nih.gov/sra). 10X PBMC, 68K PBMC, and Jurkat datasets are obtained from the website of 10X genomics (https://www.10xgenomics.com/). The worm neuron cells dataset Cao is sampled from a dataset obtained from the sci-RNA-seq platform (single-cell combinatorial indexing RNA sequencing) (http://atlas.gs.washington.edu/worm-rna/docs/). The preprocessed human tonsil data, named Tonsil, and Crohn data are available from https://portals.broadinstitute.org/single_cell. The mouse retina data and B_ lymphoma data are available from https://github.com/OSU-BMBL/marsgt.

## Code availability

scCAD is implemented in Python and the source code has been deposited at https://github.com/xuyp-csu/scCAD.

## Acknowledgments

This work was supported in part by the National Key Research and Development Program of China (No.2021YFF1201200); the National Natural Science Foundation of China under Grants (Nos. 62150048, U1909208); the Open Project of Xiangjiang Laboratory (No.22XJ02002). This work was carried out in part using computing resources at the High-Performance Computing Center of Central South University.

## Author contributions

J.X.W. and H.D.L. conceived and designed this project. Y.P.X., J.X.W., and H.D.L. conceived, designed, and implemented the scCAD. Y.P.X. and S.K.W collected datasets and conducted experiments. Y.P.X., S.K.W and H.D.L. performed the analysis. Y.P.X., Y.H.L., Q.L.F, and J.X.W. wrote the paper. All authors have read and approved the final version of this paper.

